# Probabilistic 3D-modelling of genomes and genomic domains by integrating high-throughput imaging and Hi-C using machine learning

**DOI:** 10.1101/2022.09.19.508575

**Authors:** David Castillo, Julen Mendieta-Esteban, Marc A. Marti-Renom

## Abstract

Among the existing techniques for interrogating the genome structure, Hi-C assays have become the most performed experiments and constitute the majority of the publicly available datasets. As a result, there is a continuous demand to create and improve algorithms and methods to assist the scientific community in the interpretation of Hi-C experimental data. Here we introduce probabilistic TADbit (pTADbit), a new approach that combines Deep Learning and restraint-based modelling to infer the three-dimensional (3D) structure of genome and genomic domains interrogated by Hi-C experiments. pTADbit uses thousands of microscopy-based distances between genomic loci to train a neural network model that aims at predicting the population distribution of the spatial distance between two genomic loci based solely on their Hi-C interaction frequency. pTADbit produces more accurate chromatin models compared to the original TADbit as well as other available 3D modeling methods, while drastically reducing the required computation time. The resulting ensemble of models not only agree consistently with independent measures obtained by imaging experiments but also better capture the heterogeneity of the cell population. The development of pTADbit lays the basis for the integration of data produced from high-throughput imaging assays into the 3D modelling genomes and genomic domains.

## Introduction

There are now clear evidences of the gene regulatory roles of the three-dimensional (3D) folding of the DNA inside the nucleus [1–3], many of them unveiled by Chromosome Conformation Capture (3C)-based technologies developed already twenty years ago [4]. Together with the proliferation of the 3C techniques, there has been a parallel development of methods for the inference of the 3D structure of the genome. Examples of recent developments include the reconstruction of structural models from sparse interaction data [5], from single-cell Hi-C information [6], or genome-wide low-resolution models of diploid genomes [7]. Thanks to the developed methods and their resulting models, we are gaining key insights on specific biological processes in the field of structural genomics [8]. The long list of implemented algorithms [9, 10] is a prove of the effort of the scientific community to provide the needed tools to analyze and interpret 3C-based experiments.

Among the 3C techniques, Hi-C experiments are the most applied 3C assays resulting in the majority of the publicly available datasets [11]. The main output of a Hi-C experiment is an interaction matrix representing the frequency at which two regions (or loci) of the genome are found crosslinked together within the nucleus in thousands to millions of cells. This population-based interaction matrix provides fundamental information of the 3D structure of the genome but it is, at the same time, difficult to interpret and integrate with other biological evidences as it is not a direct measure of the spatial distances at which interactions occur [12]. Therefore, an accurate measure or estimation of the physical distances at which genomic regions interact is essential for accurately characterizing how nuclear processes occur.

The inference of the 3D structure of genomes based on experiments is a process that is referred as 3D modelling. One of the strategies for the determination of those 3D conformations is restraint-based modelling in which the interaction frequency between fragments of DNA is transformed into a set of spatial restraints that are then satisfied in the resulting structures [13]. In general, finding the optimal equivalence between the frequencies and the physical distances is a key step, and requires either computationally intensive algorithms or the introduction of empirical parameters. In the first case, it is often found that the imposed restraints are similarly satisfied in conformations at different scales. In the second, the inclusion of empirical parameters might introduce some degree of arbitrary in the results. One of the available restraint-based solutions for the modelling of Hi-C information is TADbit [14], a complete Python library covering all steps in the analysis of 3C-based data. TADbit models have already provided significant biological insights (see for example [15–17]). The modeling step of the TADbit pipeline consists in the building of 3D ensembles from Hi-C interaction matrices. The conversion of interaction frequencies to physical distances involves a comprehensive and computationally expensive search of the optimal parameters that will produce models having the proper scale. Moreover, and importantly, the resulting ensemble of models, which is based on the population average contact frequency, has been shown not to fully reflect the variability observed in the cell population [18]. This inability to completely reproduce the cell-to-cell heterogeneity, far from being exclusive to TADbit, is a common drawback in many 3D modelling approaches.

Here we present *probabilistic* TADbit (pTADbit), developed to overcome the mentioned limitations by using Machine Learning (ML) in the modelling of three-dimensional genomic regions. The main idea of the new method is to use the abundant information of the recent high-throughput imaging datasets [19, 20] to produce more accurate chromatin models. Classically, measures from imaging assays like Fluorescent In Situ Hybridization (FISH) have been exclusively used to validate the accuracy of the resulting models. Nowadays, and thanks to the recent advances in the imaging of the genome [19–21], the amount of available large-scale datasets, both in terms of interrogated loci and number of cells imaged, has exponentially increased allowing for the development of predictive models based on Artificial Intelligence (AI). In pTADbit, the distances between genomic loci obtained from imaging experiments are used to train Neural Networks (NN). Such tens of thousands of image-based distances between two particular loci are sufficient to generate a smooth histogram, which can be then fitted to reconstruct a given mathematical function. Next, the NNs are trained to predict the necessary parameters to reconstruct a probability density function of distances solely from the Hi-C matrix and the genomic distance of the interacting loci (**Fig. 1** and **Methods**). Convolutional Neural Networks (CNN) are widely used in image classification and recognition tasks due to its efficient use of two-dimensional convolutional layers. pTADbit benefits from that efficiency for the extraction of features and the recognition of patterns in the Hi-C matrix, but instead of using the CNN for the classification of the images, it combines the feature extraction with a regression layer to predict a set of parameters. pTADbit results in more accurate 3D models of genomes and genomic domains when compared with the original TADbit and other available methods. It also results in ensemble of conformations with variability closer to that observed by imaging. Finally, pTADbit reduces the computation time attained by the original TADbit and other restraint-based methods.

**Figure 1.**
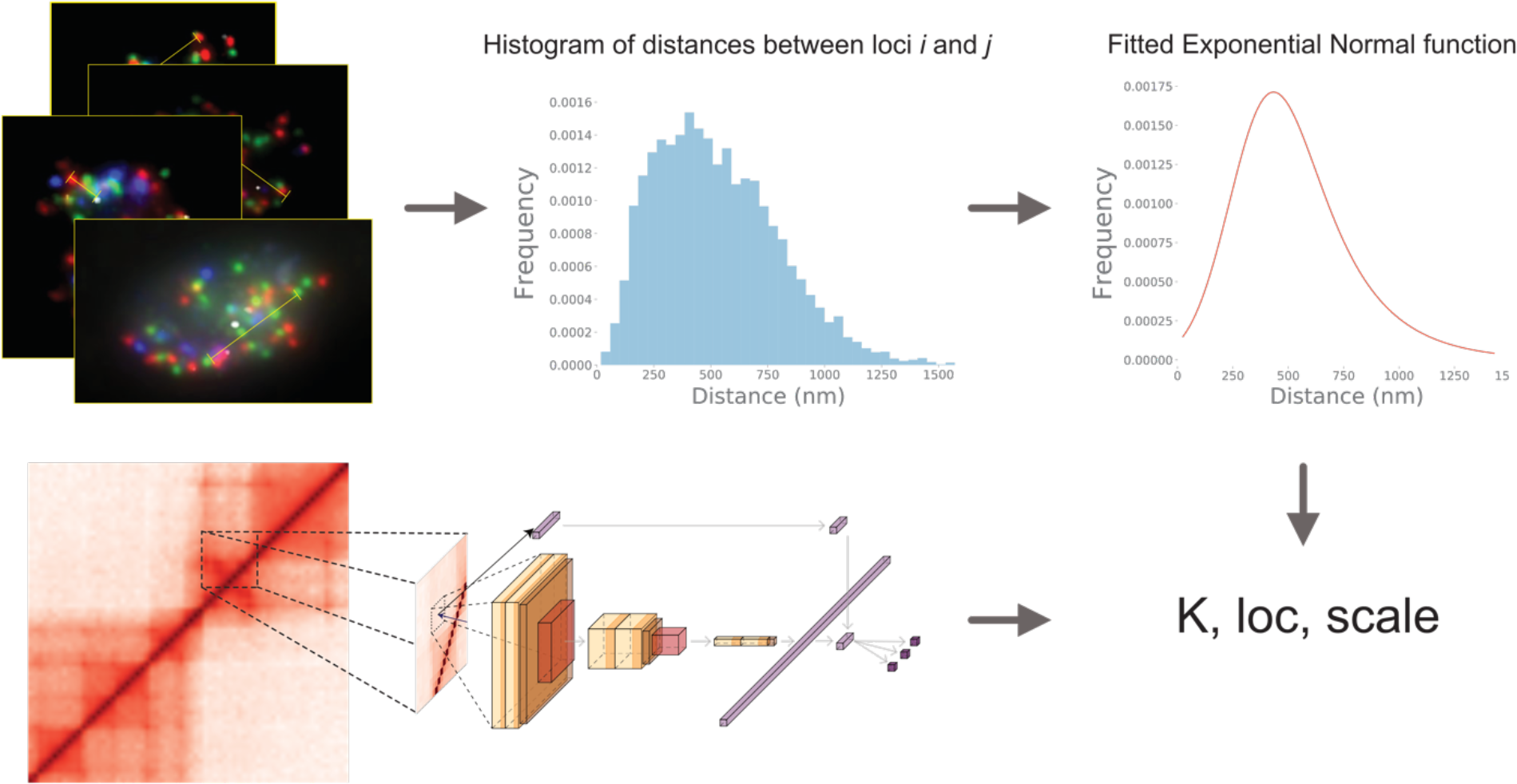
Schema of pTADbit prediction of histograms. Imaging distances are compiled into histograms which are approximated by exponential Gaussian functions. *K*, *loc* and *scale* parameters depicting the functions are predicted by Neural Networks using Hi-C matrices.

## Results

### Neural Network validation

The distances from the public imaging datasets were grouped into histograms, which were next approximated by exponentially modified Gaussian functions. Each function was depicted using three parameters: *K, loc and scale*. The goal of the trained Neural Networks (NN) was thus to predict those three parameters using only as input Hi-C data and the genomic distance between the pairs of loci which distance needs to be predicted (**Methods**). Distances between regions of 30Kbp and 250Kbp were used to train the short-range NN and the long-range NN, respectively. It is important to note that the histograms could be better approximated by other existing mathematical functions or higher-order fitting expressions like the *Beta* or a mixture of gaussian curves, but those approximations required the estimation of more variables to reconstruct the histograms. For example, the unnormalized *Beta* function is defined by four parameters compared to the three of the exponentially modified Gaussian function. Nevertheless, the error incurred in the approximation of the histograms did not significantly differ from the one obtained using other higher-order functions (**Fig. 2a**). Thus, we focused on developing NN to predict the three parameters for exponentially modified Gaussian functions.

**Figure 2.**
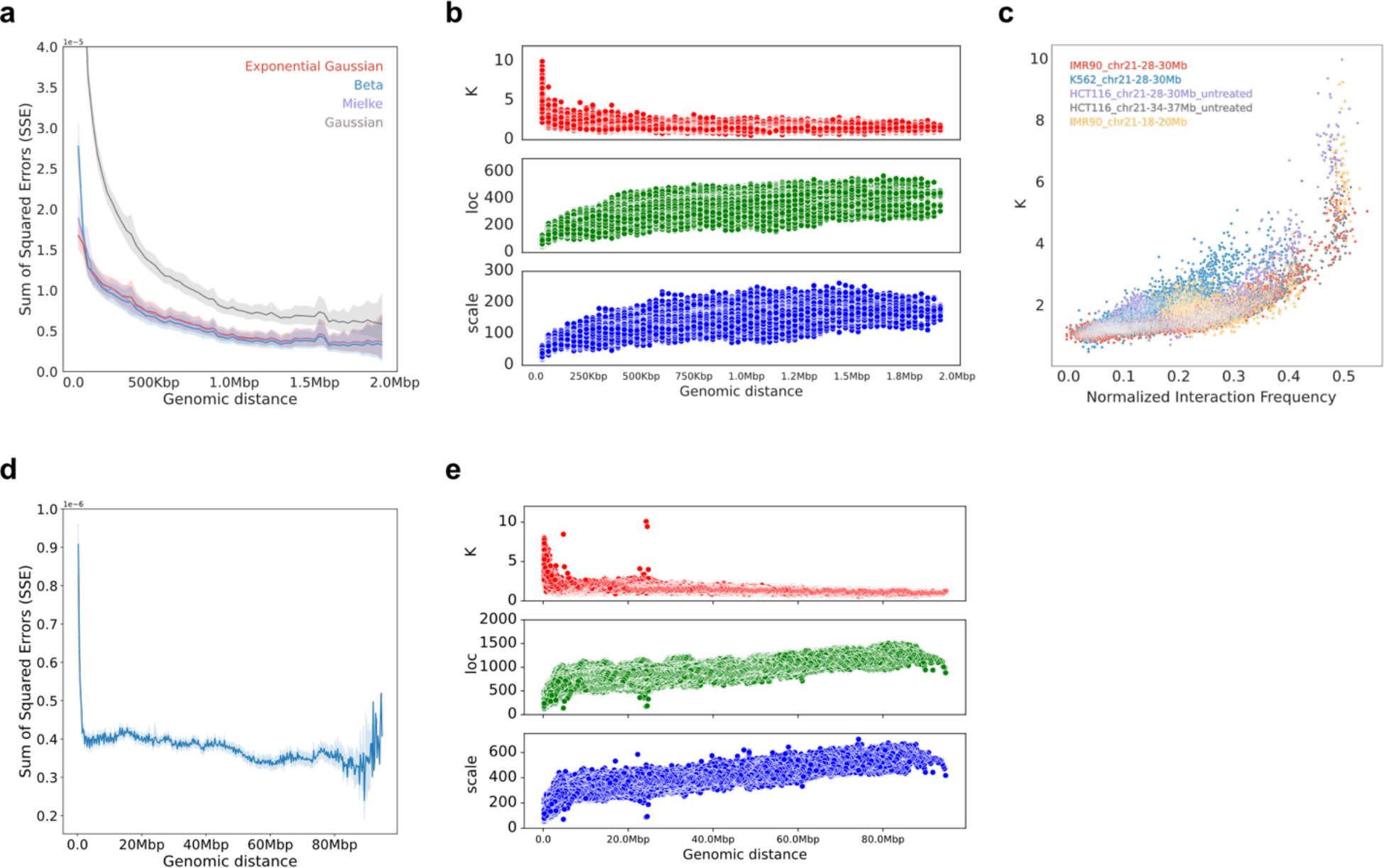
Histogram fitting. **a** Average Sum of Square Errors (SSE) of the fitting of the histograms of distances between pairs of 30Kbp regions to four mathematical curves as a function of the genomic distance. **b** *K*, *loc* and *scale* values of the fitted histograms of distances between pairs of 30Kbp regions to exponential Gaussian curves as a function of the genomic distance. **c** *K* values of the fitted histograms of distances between pairs of 30Kbp regions to exponential Gaussian curves as a function of the interaction frequency. **d** Average SSE of the fitting of the histograms of distances between pairs of 30Kbp regions to exponential Gaussian curves as a function of the genomic distance. **e** *K*, *loc* and *scale* values of the fitted histograms of distances between pairs of 250Kbp regions to exponential Gaussian curves as a function of the genomic distance.

The error in the estimation of the histograms to an exponentially modified Gaussian increased when pairs of loci were closer in genomic distance as those histograms adopt shapes that better approximate to a decreasing exponential than to a normal function (**Fig. 2a**). Additionally, the fitting of the histograms to exponential Gaussian functions resulted on a large *K* variability for pairs of consecutive loci (**Fig. 2b**). Finally, the obtained *K* values as a function of the Hi-C normalized interaction frequency (**Fig. 2c**) indicated that pairs of loci interacting with similar frequencies could be approximated by functions which *K* value considerably differed (**Fig. 2c**). Similar to the histograms of distances between 30Kbp region, we approximated the histograms of distances between 250Kbp regions with exponential Gaussian functions (**Fig. 2d**). The SSE error incurred in the approximation was considerably smaller and more constant than the obtained in the short-range histograms. We also observed a higher SSE in short genomic distances (**Fig. 2d**), which translated in a lower accuracy in the prediction of the histograms in that regime. A large variability of the *K* values in short genomic distances was also observed in the fitting of the histograms of distances between 250kbp regions (**Fig. 2e**).

Next, after determining the best function type to approximate the observed image distances, the short-range NN was trained with 11,723 different histograms from 33,755 imaged cells using 70% of the input data as a training set and the remaining 30% as a validation set. The predicted parameters *K, loc and scale* of the histograms were compared to the expected values, which resulted in high correlation coefficients (that is, *K* with *r*=0.96, *loc* with *r*=0.99, and *scale* with *r*=0.98, **Fig. 3a**).

**Figure 3.**
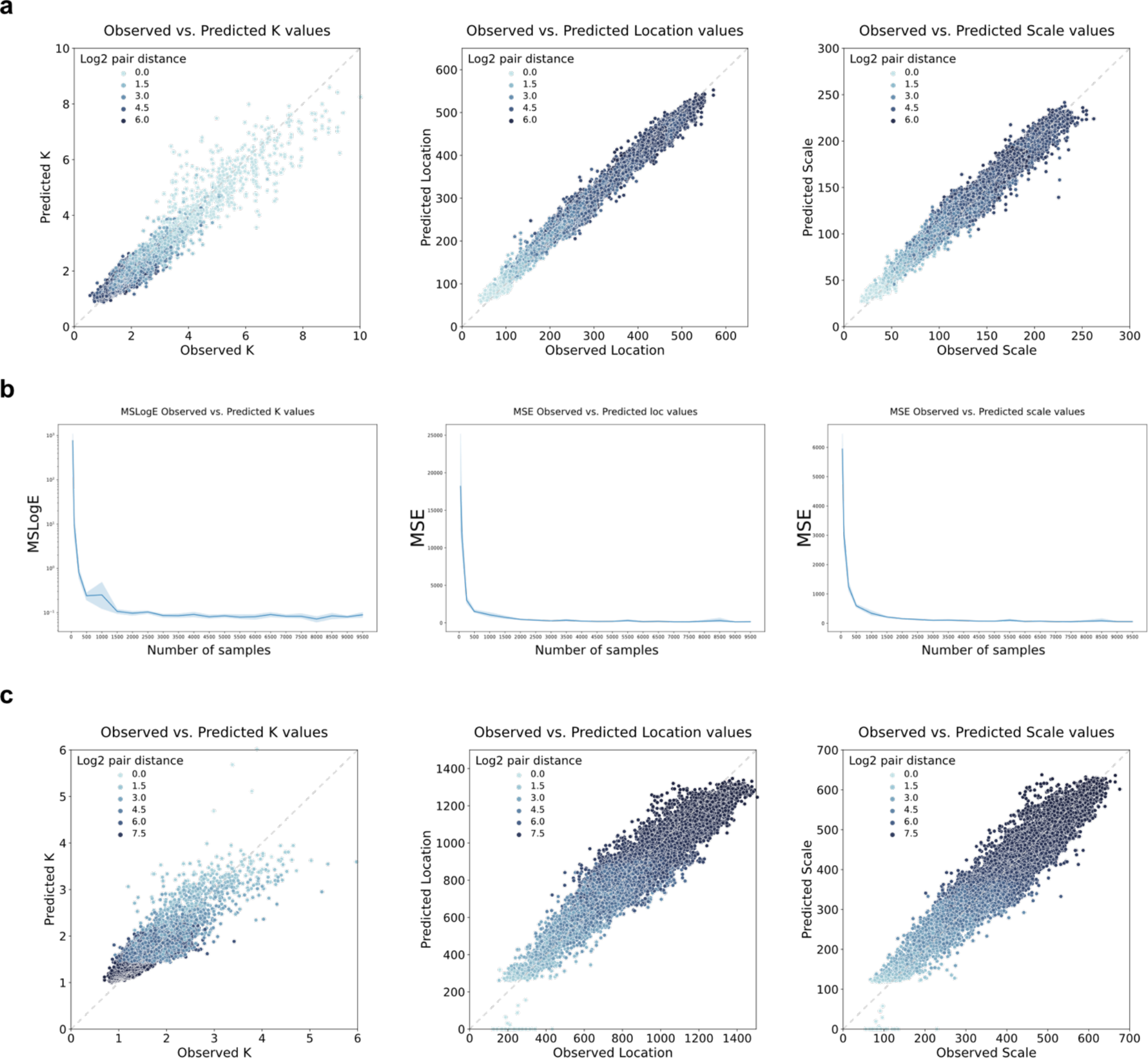
Observed vs. predicted values. **a** *K*, *loc* and *scale* values for the validation set of the short-range CNN composed by 19,298 predictions. The Pearson correlation coefficients are 0.96, 0.99 and 0.98 for *K, loc* and *scale*, respectively. **b** Mean Square Log Error of the observed vs. predicted *K* values and Mean Square Error of the observed vs. predicted *loc* and *scale* values for the validation set of the short-range CNN using an increasing number of imaged cells. The training of the CNN was repeated 5 times for each number of cells using random training and validation sets. **c** Observed vs. predicted *K, loc* and *scale* values for the validation set of the long-range NN composed by 111,669 predictions. The Pearson correlation coefficients (r) are 0.65, 0.93 and 0.93 for *K, loc* and *scale*, respectively.

Importantly, the trained NN for short-range distances resulted in very good agreement for the *loc* and *scale* parameter in all genomic distances tested. However, the agreement was not as good for the *K* value, particularly in short genomic distances characterized by high interaction frequencies, which probably reflects the above discussed inaccuracy of the histogram approximation for pairs of consecutive loci.

Next, to assess the minimum number of imaged distances (or cells) required for a good accuracy prediction by the NN, the CNNs were retrained and tested with increasing sample sizes of randomly selected cells from the *K562_chr21-28-30Mbp* dataset, which is the one set with the largest number of imaged cells (13,997 cells in total). The NN resulted in a plateau Mean Square Error around 1,500-2,000 imaged cells for the three measures of *K*, *loc*, and *scale* (**Fig. 3b**). Interestingly, this is a similar number of cells required to obtain dense Hi-C interaction maps using the so-called “low-input” protocols [22], which may indicate that this is the minimum number of cells required to properly capture the variability in genome structure in a population.

Finally, the long-range NN was trained with 61,789 histograms from 4,848 imaged cells using 70% of the input data as training set and the remaining 30% as validation set. The predictions resulted in high correlation coefficients with the expected values for each parameter *K* (r=0.87), *loc* (r=0.94) and *scale* (r=0.93) (**Fig. 3c**).

### pTADbit benchmarking

Numerous algorithms for the 3D modelling of chromatin exist [9, 13]. It is, however, difficult to find implemented methods publicly available that can be directly compared with pTADbit. Many of the existing packages follow a different approach by providing unique consensus solutions instead of ensembles of structures. Others are tailored to model the genome at lower resolutions and are simply not easy to adapt to building structures at 30Kbp. Despite these limitations, together with the original TADbit method [14], we executed the Lorentzian 3D Genome (LorDG) [23] and the Chrom3D [24] packages and compared their resulting models to those obtained by pTADbit. The three methods, as for pTADbit, are able to provide ensembles of structures at the high-resolution (30Kbp) using as input solely the Hi-C interaction matrix. Next, for the three methods, we generated an ensemble of 1,000 models of the region 40Mbp-42.5Mbp in chromosome 21 in IMR90 using as input a Hi-C interaction matrix (**Fig. 4a-d**). As this genomic region has also been imaged but never used for the NN training, we were able to directly compare the results of the models against observed image distances.

**Figure 4.**
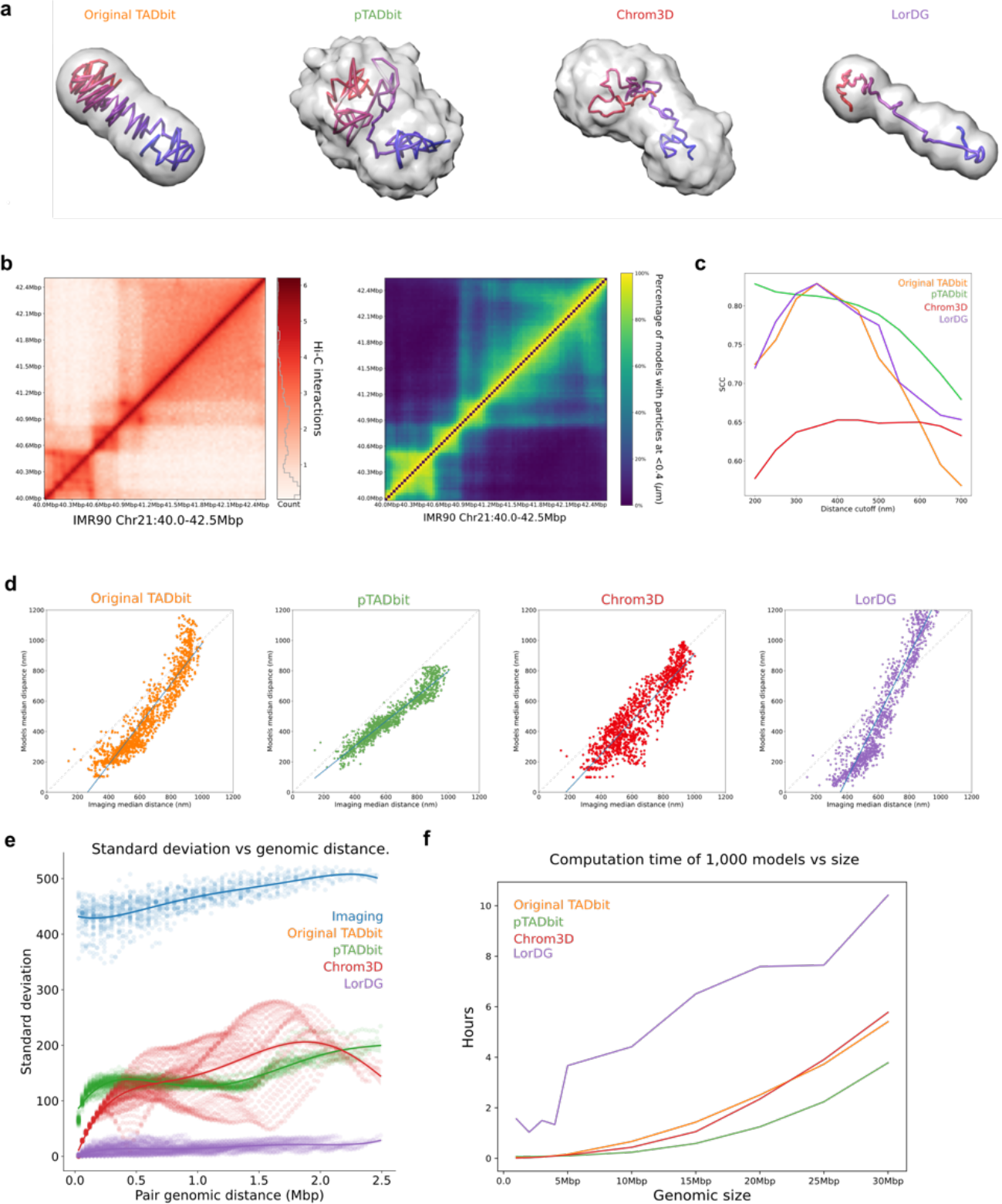
Model benchmarking. **a** Ensemble of models at 30Kbp of the genomic region 28Mbp-30Mbp in chromosome 21 in IMR90 generated using TADbit (orange), pTADbit (green), Chrom3D (red) and LorDG (purple). The centroid model (the one closer to the average) is depicted by a tubular shape colored from blue (40Mbp) to red (42.5Mbp) and the occupancy of the ensemble is represented by a semi-transparent grey shadow. **b** Normalized Hi-C matrix of the genomic region used to produce the ensembles in **a** (left panel) and contact map of the pTADbit ensemble using a distance cutoff of 400 nm (right panel). **c** Stratum-adjusted Correlation Coefficient (SCC) between the contact map of the ensembles in **a** and the normalized Hi-C matrix in **b** using different distance cutoffs and for all compared methods. **d** Comparison of the pairwise median distances between each 30Kbp probe in the imaged genomic region 40Mbp-42.5Mbp in chromosome 21 in IMR90 (n=7,591 cells) and all loci in the ensemble of 1,000 models of the same region produced by the original TADbit (orange), pTADbit (green), Chrom3D (red), and LorDG (purple). The Pearson correlation coefficients (r) were 0.92 for TADbit, 0.96 for pTADbit, 0.9 for Chrom3D and 0.95 for LorDG. **e** Standard deviation of the distance between pairs of loci depending on its genomic distance in the ensemble of 1,000 models. **f** Computation time to produce an ensemble of 1,000 models of different genomic sizes for the modelled region. Results were obtained using 24 cores in a workstation with an Intel(R) Core (TM) i9-7960X @ 2.80GHz with 128 Gb of RAM.

An indirect way of benchmarking the generated models is to assess the agreement of a contact map calculated from the generated ensemble of 3D confirmation with that of the input Hi-C interaction matrix [14]. This is accomplished by producing contact maps simulating the crosslinking in the models at different cutoff distances. For example, the comparison of the 30Kbp resolution contact map of the genomic region chr21:28-30Mbp in IMR90 obtained with pTADbit with 400nm cutoff distance (**Fig. 4b**) results in a Stratum-adjusted Correlation Coefficient (SCC), a metric designed specifically to compare Hi-C matrices [25], of 0.81. All the ensemble of models built by the compared methods result in high SCC values for cutoff distances below 400nm except Chrom3D with values that are slightly lower than the other ensembles. pTADbit and Chrom3D results are more consistent across all distance cutoffs until 500nm (**Fig. 4c**). Interestingly, the original TADbit has a level of accuracy similar to pTADbit for the short range contacts as TADbit indeed optimizes the most likely distance of an interaction given the input matrix [14]. However, the original TADbit clearly suffered in identifying longer-range interactions present in the input Hi-C. This is not the case of the results from pTADbit.

To have a more direct benchmarking of the models, we turned to independent imaging experiments never used for modeling. We compared the pairwise median distances between all loci in the models with the pairwise median distances of high-throughput images from the literature (**Fig. 4d**) [20]. As LorDG and Chrom3D do not explicitly have a scaling factor to a priory assess the real size of the resulting ensemble of models, we adjusted their size multiplying the model coordinates by the median distances of the pTADbit ensemble. That is, LorDG model distances were multiplied by 92.19 and Chrom3D by 49.49. The scaling of the LorDG and Chrom3D ensembles simplified the comparison with the other ensembles. All methods resulted in good correlations for all modeled cases (r=0.92 for TADbit, r=0.96 for pTADbit, 0.90 for Chrom3D, and 0.95 for LorDG, and **Fig. 4d**) but the match of the distances with the ones obtained in the images differed considerably. The original TADbit ensemble exhibit a transformation that undervalued short distances and did not grow linearly. The median distances in the pTADbit ensemble of models grew linearly at almost the same rate as in the images but with a slightly scale offset. This scale offset could be caused by differences in the protocols or conditions used in the acquisition of the images in this dataset compared to the ones used in the datasets of the training of the NNs. Chrom3D did not consistently reproduce the distances as the algorithm emphasizes a subset of bead pairs that significantly interact instead of optimizing distances between large number of pairs. Finally, LorDG transformation was linear but it also undervalued short distances and overvalued the long ones.

Next, to assess if the models reproduce the distance variability observed in the images, we plotted the standard deviation of the pairwise distances with the genomic distance obtained in the ensembles (**Fig. 4e**). We observed that TADbit and LorDG models were too deterministic for all ranges of genomic distances. That is, did not result in an ensemble that captured the variability observed in the images. In turn, Chrom3D models resulted also in lower variability in very short genomic distances. This was likely a consequence of constraining only pairs which interaction values are statistically significant. pTADbit ensemble resulted in a constant distance variability with the exception of a decrease in variability for very short genomic distances. Although pTADbit and Chrom3D exhibit an increase of the variability of the ensemble of solutions observed in imaging experiments, such variability is still lower for the generated models. The reduced variability of distances in the solutions is further addressed in the **Discussion** section.

Finally, the computational burden of generating an ensemble of 1,000 models using the four methods was assessed on a computational workstation with a 24 core Intel(R) Core (TM) i9-7960X @2.80GHz with 128 Gb of RAM. The ensemble of models was generated for models of increasing size between 1Mbp and 30Mbp (**Fig. 4f**). All methods followed an exponential increase trend as the size of models increase with the exception of LorDG, which appeared to follow a more linear trend. However, LorDG compared worse against all other methods in all tested genomic sizes. pTADbit, in contrast, favorably compared against all other methods for models larger than 7-8Mb, with a reduction of computational time larger as the size of the models increased. In average, pTADbit required about 3.5h of computational time to generate 1,000 models of 30Mbp of size at 30Kbp resolution (that is, 1,000 particles), which is about two thirds the time required for TADbit or Chrom3D.

### Modeling additional regions

After validating the ability of pTADbit to recover regions of the genome used in the training phase, we next generated with pTADbit an ensemble of 1,000 structural models for each of the regions 54Mbp-58Mbp and 151Mbp-155Mbp of chromosome 4 in human foreskin fibroblasts (HFFs). The contact maps obtained from the ensembles (**Fig. 5a,b**) resulted in high correlation with their equivalent Hi-C matrices (SCC at 400nm of 0.60 and 0.81, respectively). The median distances between various particles in the ensembles of models with the distances of publicly available imaging data [26] labelling a total of 18 regions with bacterial artificial chromosome (BAC) probes also resulted in high correlations of r=0.93 (**Fig. 5c**) and r=0.89 (**Fig. 5d**) for the 54Mbp-58Mbp and 151Mbp-155Mbp in chromosome 4 in HFF, respectively. Finally, we verified that the contacts maps of the ensembles agreed with the published measures at different distance cutoffs (150, 200 and 350 nm, **Fig. 5e**). The contact maps resulted in high correlation with the published percentages for each of the cutoffs (r=0.83, 0.84 and 0.86 respectively), confirming the accuracy of the obtained models.

**Figure 5.**
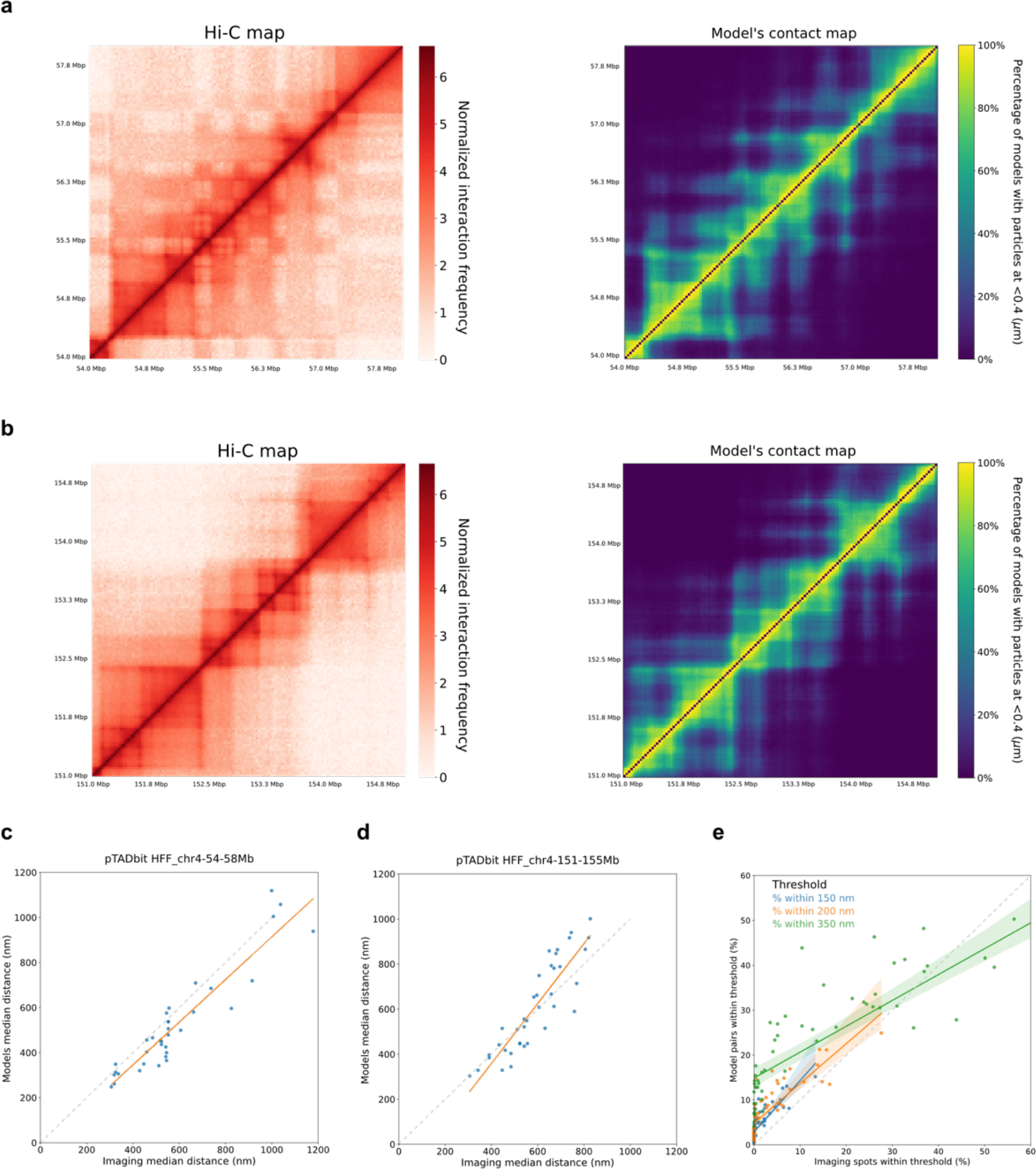
Model accuracy. **a** Hi-C matrix of the genomic region 54Mbp-58Mbp in chromosome 4 in HFF (left) and contact map (right) using a distance cutoff of 400 nm of the ensemble of 1,000 models generated from the matrix in a using pTADbit. **b** Hi-C matrix of the genomic region 151Mbp-155Mbp in chromosome 4 in HFF (left) and contact map (right) using a distance cutoff of 400 nm of the ensemble of 1,000 models generated from the matrix in a using pTADbit. **c** Comparison of the pairwise median distances between each of the 9 probes in the imaged genomic region 54Mbp-58Mbp in chromosome 4 in HFF (n=50,197 distances) and the corresponding loci in the ensemble of 1,000 models of the same region produced by pTADbit. The Pearson correlation coefficient (r) is 0.93. **d** Same as **b** for the genomic region 151Mbp-155Mbp (n=116,047 distances) resulting in a Pearson correlation coefficient (r) of 0.89. **e** Comparison of the percentages of pairs of the 9 imaged probes which distance is within 150, 200 and 350 nm each in the imaged genomic regions 54Mbp-58Mbp and 151Mbp-155Mbp in chromosome 4 in HFF and the corresponding loci in the ensemble of 1,000 models of the same region produced by pTADbit. The Pearson correlation coefficients (r) are 0.83, 0.84 and 0.86 for 150, 200 and 350 nm, respectively.

### Modelling full chromosome 19

With the reduction of the computation times in the generation of the ensembles, pTADbit can now be applied to model large regions of the genome at high resolutions, including entire chromosomes.

We next modeled the entire human chromosome 19 at 30Kbp (**Fig. 6a**), which results in a total of 1,971 particles. The ensemble of 1,000 models required a total of about 30h of computational time on a single workstation with a 24 core Intel(R) Core (TM) i9-7960X @2.80GHz with 128 Gb of RAM. As in the previous validations, the contact map of the ensemble (**Fig. 6b**) resulted in high correlation with the Hi-C matrix of the chromosome (SCC=0.79). Finally, the standard deviation of the pairwise distances across different genomic distances indicated that pTADbit may not capture full variability for very short distances while resulting in larger standard deviations for larger ones (**Fig. 6c**). This was likely the result of combining the modelling of low- and high-resolution structures where the Monte Carlo simulations of the low-resolution models are twice as the ones used in the high-resolution ones (**Methods**).

**Figure 6.**
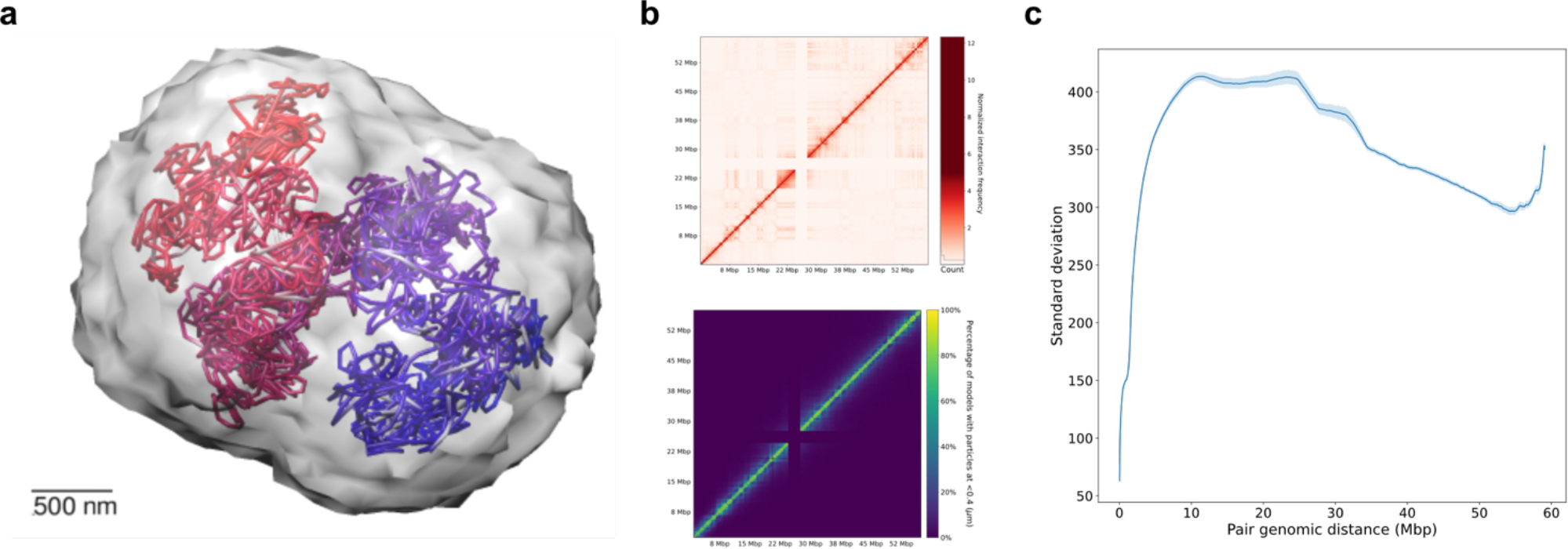
Model accuracy for an entire chromosome. **a** Ensemble of models at 30Kbp of the chromosome 19 in IMR90 generated using pTADbit. The centroid model (the one closer to the average) is depicted by a tubular shape colored from blue to red and the occupancy of the ensemble is represented by a semi-transparent grey shadow. **b** Normalized Hi-C matrix of the chromosome 19 used to produce the ensemble of models (top panel) and contact map of the pTADbit ensemble using a distance cutoff of 400 nm (lower panel). **c** Standard deviation of the distance between pairs of loci depending on its genomic distance in the ensemble of 1,000 models.

## Discussion

pTADbit provides an update of the original TADbit [14], which makes use for the first time of large-scale image data to train the required transformation of frequency of interactions observed using Hi-C and the physical distance between loci measured by imaging technologies. Indeed, this transformation is key to any algorithm aiming at modeling genomes and genomic domains. Briefly, for the methods benchmarked in this study, the original TADbit included a configurable scale factor that sets the amount of DNA in base-pairs contained in one nanometer. The scale factor drives the structures to the proper range of distances during the optimization step in which a grid search is conducted to find the optimal transformation. Chrom3D relies on two configurable parameters to produce the structures with the appropriate scale: the nuclear radius and the volume of the modelled chromosome as a percentage of it. It does not include any scaling factor for the modelling of genomic regions shorter than full chromosomes. LorDG optimizes ∝ in the transformation:

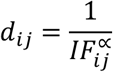

where *d*_*ij*_ is the distance between particles *i* and *j* and *IF*_*ij*_ its interaction frequency but it does not include any scaling factor. The new pTADbit is thus the first method to implicitly incorporate this transformation in the NNs by integrating the training imaging datasets. The explicit transformation from Hi-C data to distances is one the strengths of pTADbit compared to existing methods because it allows the reduction of the computation times in the generation of the ensembles.

The approach to incorporate the imaging information in the modelling method consists in the grouping of single cell distances into population-based histograms in which the information of the exact conformation adopted by chromatin in each individual cell is lost. Those histograms are then predicted and used in the production of ensembles of individual structures that follow the probabilistic distributions. The impossibility of the prediction of the precise histogram solely from the Hi-C information is overcome by their approximation to exponential Gaussian curves, which can be depicted using three parameters *K*, *loc* and *scale* that can be accurately predicted by the NNs. The approximation of the histograms using other higher-order curves decreased the error of the fitting particularly for distances between consecutive regions but made their prediction more computationally intensive as it required the optimization of additional parameters. Importantly, the accuracy of the prediction for distances between consecutive loci did not significantly change the shape of the chromatin structures, which is mainly driven by distances between far apart regions.

The limited number of high-resolution high-throughput imaging datasets is a restricting factor for the size of the modelled regions. The distances used in the short-range NN were obtained from traces of chromatin regions of 2Mbp length at 30Kbp. Therefore, the histograms predicted by the short-range NN are, conceptually, not valid for pairs of fragments that are farther than the 2Mbp, which restricts the total length of the modelled region. To overcome this limitation pTADbit implements a multi-resolution approach where distances from traces at lower resolution (250Kbp) are used to train the long-range NN. Then, in the modelling process, low-resolution models are built first to determine the shape of the structures at genomic distances that are above the limitation imposed by the short-range NN. The distances of those low-resolution models are then used to produce the final high-resolution conformations.

Recent studies highlight the importance of providing ensembles of structures as opposed to single consensus solutions better capturing the heterogeneity of the cell population [7]. This is indeed one of the main objectives of pTADbit. However, pTADbit benchmarking still results in variabilities smaller than those observed by microscopy (although significantly larger than other compared approaches). The main reason relies in the assignment of the restraints during the modelling approach (**Methods**). Each reconstructed histogram is used as a probability distribution function from where to sample the distance restraints. The restraints imposed are mutually independent and randomly obtained from the distributions. If, during the sampling process, a distance from the tail of the distribution (far from the median) is assigned between fragment *i* and *j*, the probability of having a similar distance assigned to *i* and the neighboring fragments of *j* is low. As a result, the obtained structures, although having a high degree of variability, tend to penalize structures which pairs of distances have very low probability to occur in the population.

We demonstrate that the ensemble of structures obtained with pTADbit not only are in high agreement with both Hi-C interaction matrices and independent imaging data but they are also closer to represent the large heterogeneity of the cell population. In that respect, we could have further increased the degree of variability observed in the imaging data by increasing the number of Monte Carlo simulations and keeping only the structures that have best satisfied the imposed restraints. However, that would have increased the computation time, which is another of the advantages of pTADbit. Moreover, although having imaging data from only a few regions of the genome, the method is applicable to the rest of the human genome. However, we expect small biases in the distance predictions caused by the scarcity of datasets used for the training of the NNs. Indeed, only imaging data of specific regions of chromosome 21 are available at high-resolution. We anticipate that the release of new high-throughput imaging datasets of different regions will allow us to increase the accuracy of the predictions. Moreover, pTADbit is not restricted to the existing trained NNs and it is prepared to use other future TensorFlow networks trained with more extensive datasets.

In summary, pTADbit is a novel approach for the modelling of chromatin fiber that makes use of imaging information to accelerate the generation of ensembles of 3D structures and to reproduce more accurate models that better capture heterogeneity of the cell population.

## Methods

### pTADbit architecture

TADbit, a broader pipeline for the analysis of Hi-C data, from the mapping of the sequenced reads to the production of 3D ensemble, now includes pTADbit as part of its python package. Although pTADbit represents a completely different approach in the generation of ensemble of genomic regions, TADbit is the perfect container providing the required tools to produce the Hi-C matrices and analyze the resulting structures. The neural networks used in pTADbit were built using TensorFlow [27] and can be replaced by other TensorFlow models as far as input and output parameters are conserved.

### Prediction of distance distributions

A short-range Convolutional Neural Network (CNN) and a long-range Neural Network (NN) were trained to predict the distribution of distances between two chromatin loci from its Hi-C interaction frequency and neighborhood. The short-range CNN was trained to predict distributions in Hi-C matrices at 30Kbp resolution and can be used to determine the structure of the models at the finer scale of TADs and sub-TADs (shorter than 1.5Mb). The long-range NN predicts distributions using Hi-C matrices of 250Kbp and is used to shape the models in genomic ranges that are usually larger than the average size of mammalian TADs (larger than 1.5Mb).

### Histogram fitting

The short-range CNN was trained to predict histograms of distances obtained from extensive imaging between genomic loci *i and j* using as input the interaction frequency, the genomic distance and the corresponding Hi-C sub-matrices centered at *ij*. The long-range NN, instead, only used the interaction frequency and the genomic distance as input. Distances were obtained from publicly available high-throughput datasets of labelled regions in individual cells [19, 20].

For each cell, coordinates of the different imaged loci were converted to pairwise distances between loci *i and j*. Next, the distribution of the distances *ij* imaged in the same cell type and genomic region were stacked for 200 bins as “Observed data” (**Fig. 1** second panel). Next, an exponentially modified Gaussian distribution was fit to each histogram (**Fig. 1** third panel). The probability density function of the exponentially modified Gaussian distribution was:

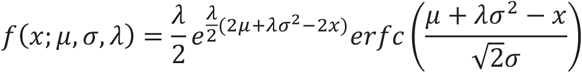

where *erfc* is the complementary error function defined as:

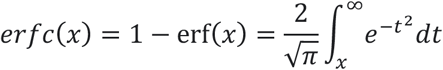

Such fitting was used as implemented in the *SciPy* python package (https://scipy.org). The three parameters *K, loc and scale* in the parameterization thus corresponded to having *loc* and *scale* equal to *σ* and *μ*, respectively, and *K*=1/(*σλ*). The objective of this procedure was to represent and be able to reconstruct the histograms using the fewer number of parameters as possible; in our case the triad *K*, *loc* and *scale*, which in turn will be predicted by our trained NNs.

### Short-range Convolutional Neural Network

The short-range CNN (**Fig. 7a**) was composed by two inputs: one encoder for the submatrix that was formed by three convolutional and two pooling layers and another encoder for the genomic distance with a fully-connected layer. They both converged to a fully-connected layer with three outputs *K*, *loc* and *scale*. Weights were trained using the Adam optimizer [28] to minimize the mean-squared error (MSE) between the input and the output. Rectified linear unit (ReLU) activation functions were used for hidden layers and softplus functions were used for the outputs to prevent the prediction of negative values. The CNN was trained using as input datasets from Bintu et al. 2018 [19] (**Table 1)**. The list of coordinates of the centers of the imaged 30Kbp segments were obtained from https://github.com/BogdanBintu/ChromatinImaging and used to calculated the *K*, *loc* and *scale* parameters of each pairwise distance as described in the previous section. Additionally, Hi-C matrices of chromosome 21 were obtained from GEO database (GSE63525 [2] and GSE104334[29]), normalized using Vanilla coverage [30] and scaled to the range 0 to 1. Then, for each pair of genomic loci *i and j*, its *K*, *loc* and *scale* parameters were matched to the Hi-C 19×19 submatrix centered in the intersection of *ij*. Together with the genomic distance between loci *i and j*, the Hi-C submatrix was used as input data to train the CNN. In addition, each *K*, *loc* and *scale* parameters was assigned to Hi-C matrices with different sequencing depths to assure that the predictions were not tied to a specific quality of the matrix. To evaluate the performance of the CNN, a K-Fold Cross Validation [31] with 5 folds and 3 repeats was applied and an average mean absolute error of 337.66 with a standard deviation of 18.39 was obtained. The reduced standard deviation with respect to the average indicates that the performance of the CNN did not depend on any specific partitioning of the training and validation sets and was independent of their random selection.

**Figure 7.**
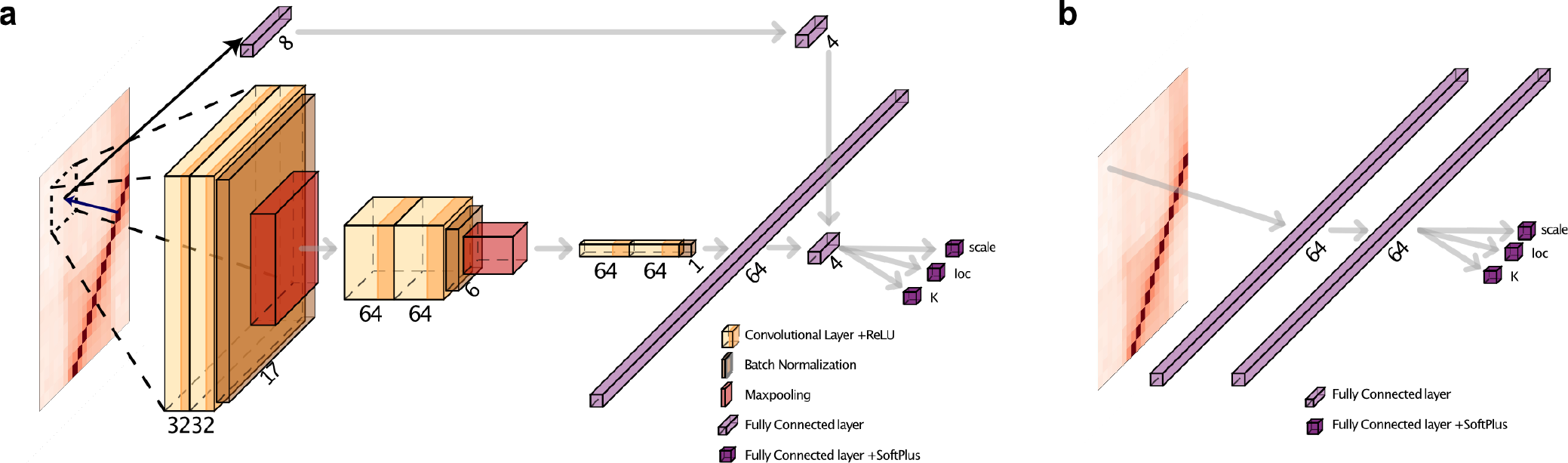
Convolutional Neural Network architectures. **a** Short-range Neural Network architecture. The 19×19 pixels input sub-matrices pass through the initial convolutional layers that performs a hierarchical decomposition of the information allowing the CNN to learn a wide range of features, from the very local to the more global ones. The subsequent fully connected layers learn non-linear combinations of the extracted features, combine them with the genomic distance of the center of the sub-matrix and does a linear regression to estimate *K*, *loc* and *scale*. **b** Long-range Neural Network architecture. The long-range NN is a simplified version of the short-range where the convolutional layers are removed and the input consists only in the interaction frequency and the genomic distance.

**Table 1.** Image datasets used in the different trained CNN.

| CNN | Label | Cell line | Region (hg38) | # Cells | # Pairwise distances |
| --- | --- | --- | --- | --- | --- |
| Short-range | IMR90_chr21-18-20Mb | IMR90 | 18,627,714-20,577,518 | 1,277 | 2,080 |
| Short-range | IMR90_chr21-28-30Mb | IMR90 | 28,000,071-29,949,939 | 4,871 | 2,080 |
| Short-range | K562_chr21-28-30Mb | K562 | 28,000,071-29,949,939 | 13,997 | 2,080 |
| Short-range | HCT116_chr21-28-30Mb_untreated | HCT116 | 28,000,071-29,949,939 | 1,979 | 2,080 |
| Short-range | HCT116_chr21-34-37Mb_untreated | HCT116 | 34,628,096-37,117,534 | 11,631 | 3,403 |
| Long-range | Chr2-p-arm_replicate | IMR90 | 1-94,750,001 | 4,848 | 61,789 |

### Long-range Neural Network

The long-range NN (**Fig. 7b**) was composed by two fully-connected layers with the interaction frequency as input and three outputs *K*, *loc* and *scale*. Weights were trained using the Adam optimizer to minimize the mean-squared error (MSE) between the input and output. Rectified linear unit (ReLU) activation functions were used for hidden layers and softplus functions were used for the outputs to prevent the prediction of negative values. The NN was trained with the imaging dataset of the p-arm of chromosome 2 from Su et al. 2020 [20] (**Table 1)**. Similarly, the Hi-C matrix was obtained from the GEO database (GSE104334[29]). We assigned the list of coordinates of the centers of the imaged segments to their equivalent 250Kbp bins of the Hi-C matrix and proceed similarly to the short-range CNN but in this case using just the Hi-C interaction frequency and the genomic distance of each pair of loci. Therefore, the long-range neurons are trained to perform a regression without the image recognition explained in the short-range CNN.

### Bead-on-a-string models

Each 30Kbp bin (column or row) of the input Hi-C interaction matrix was represented by a spherical particle which size was proportional to the number of nucleotides contained in the DNA fragment. Those particles form a connected chain that mimic the polymeric nature of DNA in what is commonly referred as a bead-on-a-string model.

### Assignment of spatial restraints and scoring

To combine short and long-range restraints a multi-scale approach was adopted in the modelling process by building low-resolution models and using their distances to restraint long-range interactions. To produce the ensemble of models at low resolution, the input Hi-C matrix was first downscaled to 250Kbp and used as input for the long-range NN to predict their corresponding *K*, *loc* and *scale* parameters from the input frequency and genomic distance between the interrogated loci. Using the predicted parameters of the exponentially modified Gaussian function, the distribution of distances between two particles *i and j* in the population of models was recreated. Distances were randomly sampled from the distributions and assigned as spatial restraints between *i* and *j* with the following criteria:

- Consecutive particles were always restrained to guarantee the continuity of the polymer chain. The absence of restraints between two consecutive particles might result in the disconnection of the chain, which is incompatible with the polymeric nature of chromatin.
- For non-consecutive particles, 60% of the possible pairs were randomly selected in each individual model and restraint with a distance sampled from the distributions. Selecting different restraint pairs in each structure increased the heterogeneity of the resulting ensemble. By using a small number of restraints, the computation time of the model was reduced at the same time that the variability of the resulting conformations increased. Indeed, the assignment of all possible restraints in each individual model would favor the introduction of contradictory restraints if, for example, regions were brought together to close distance while neighboring regions were taken apart. It is important to note that restraints that could potentially produce triangle inequalities were discarded during the assignment of restraints in each individual model. That is, for every triad of particles *i*, *j*, *k* the assignment of more than two distance restraints between them was prevented. We found the percentage balance 60/40 between restraint and non-restraint pairs to be a good compromise between computation time and variability of the resulting ensembles.
- The remaining 40% of the non-consecutive pairs were not restrained.

The spatial restraints used between pair of particles were implemented as harmonic oscillators that penalize quadratically deviations from the given equilibrium distance. The mathematical function of the restraint was:

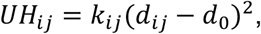

where *d_ab_* is the distance between particle *i* and particle *j* in the model, *d_0_* is the predicted equilibrium distance sampled from the NN distribution and *k_ab_* is the harmonic constant that depends on the *loc* parameter of the predicted distribution as follows:

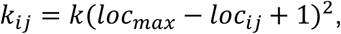

being *loc_max_* the maximum *loc* parameter predicted for the model.

The sum of all the imposed harmonic restraints forms an objective function to minimize. To reach that goal, a Monte Carlo simulated annealing sampling approach was used where the conformation of the 3D model was randomly modified and the new configuration accepted or rejected according to the Metropolis criteria [32]. The simulation was repeated with the spheres at different starting random positions, generating each one a single model. The number of structures produced were two times the number initially requested, conserving at the end of the process only the half that best satisfied the imposed restraints. By increasing the number of simulations resulted in structures that were more compatible with the imposed restraints. Considering the low-resolution of these models, increasing the number of requested structures did not compromise the computation time. Finally, the distances between pairs of particles in the low-resolution models were used to build the models at higher resolution.

Next, low-resolution models at 250Kbp were used to produce high-resolution models at 30Kbp by imposing as restraints all distances at 250Kbp for pairs of particles which genomic distance was above the applicability of our short-range NN (that is, 1.5 Mb). Briefly, each 30Kbp particle in the high-resolution model was assigned to the 250Kbp containing it as distances of the low-resolution models were considered as a good approximation to the distances of the high-resolution structures. Each low-resolution structure was then used in the production of one high-resolution structure, as to maintain the composition of the already optimized low-resolution ensemble. During the building of the high-resolution models, the Hi-C submatrices at 30Kbp resolution were used as input to the short-range CNN to predict the *K*, *loc* and *scale* parameters of the exponentially modified Gaussian functions of each pair of loci *i* and *j*, which genomic distance was below 1.5Mbp. For pairs which genomic distance was larger than 1.5Mbp, instead of sampling from the distributions, the optimized distances obtained in each of the low-resolution models were used as restraints.

Analogously to the low-resolution models, the distribution of distances between pairs of particles which genomic distance was below 1.5Mbp were recreated and randomly sampled to assign the distances as spatial restraints. The assignments were as follows:

- Consecutive particles were always restrained to guarantee the continuity of the polymer chain.
- For non-consecutive particles which genomic distance was below 1.5Mbp, 30% of the possible pairs were randomly selected in each individual model and restraint with a distance sampled from the distributions. We found that the assignment of 30% of the restraints in the short genomic regime was enough to recreate the structural features without compromising the computation times.
- For non-consecutive particles which genomic distance was above 1.5Mbp, 60% of the possible pairs were randomly selected in each individual model and restrained with the distances obtained from its starting low-resolution model.
- The remaining non-consecutive were not restrained.

Finally, the ensemble of conformations was built by minimizing the objective function with a Monte Carlo sampler.

### Three-dimensional (3D) modeling

The 3D models in pTADbit were generated by assigning the predicted distances between loci as spatial restraints that were then satisfied using the Integrative Modeling Platform (IMP) [33]. The procedure is in many ways similar to the previously published TADbit [14].

### Stratum-adjusted correlation coefficients (SCC)

To calculate the SCC coefficient, first the matrix is smoothed with a 2D mean filter to minimize the effect of noise and biases and second, the Hi-C data is stratified according to their genomic distance.

The smoothing filter is characterized by the span size *h* and the correlation is limited to a maximum number of diagonals starting from the main one. The SCC measures in this manuscript were calculated using a span size *h* of 3. In the case of the full chromosome 19, 4Mbp was used as the maximum distance from the diagonal at which SCC was computed. All the diagonals were taken into account for the rest of the matrices.

### Production of the ensemble of models with TADbit, LorDG and Chrom3D

#### TADbit

The ensemble of 1,000 models generated with the original TADbit algorithm [14] was obtained by performing an optimization step to find optimal values of −0.4 for *lowfreq*, 0 for *upfreq* and 430 nm for the distance *cutoff* using a *scale* of 0.007. With those values, the final ensemble of models was obtained as previously described [14].

#### Chrom3D

The ensemble of 1,000 models generated with the original Chrom3D algorithm [24] was obtained following the publicly available protocol in https://github.com/Chrom3D using a *cooling-rate* of 0.001. The generation of individual models was parallelized with an in-house python script that assigned a different seed number to each run.

#### LorDG

The ensemble of 1,000 models generated with the original LorDG algorithm [23] was obtained following the publicly available protocol in https://github.com/BDM-Lab/LorDG. LorDG estimated the ∝ parameter in the relation between the interaction frequency and the physical distances to be 0.6.

## Acknowledgments

MAM-R acknowledges support by the Spanish Ministerio de Ciencia e Innovación (PID2020-115696RB-I00) and the National Human Genome Research Institute of the National Institutes of Health under Award Number RM1HG011016. The content is solely the responsibility of the authors and does not necessarily represent the official views of the National Institutes of Health. CRG acknowledges support from ‘Centro de Excelencia Severo Ochoa 2013-2017’, SEV-2012-0208 and the CERCA Programme/ Generalitat de Catalunya as well as support of the Spanish Ministry of Science and Innovation through the Instituto de Salud Carlos III and the EMBL partnership, the Generalitat de Catalunya through Departament de Salut and Departament d’Empresa i Coneixement, and the Co-financing with funds from the European Regional Development Fund (ERDF) by the Spanish Ministry of Science and Innovation corresponding to the Programa Operativo FEDER Plurirregional de España (POPE) 2014-2020 and by the Secretaria d’Universitats i Recerca, Departament d’Empresa i Coneixement of the Generalitat de Catalunya corresponding to the programa Operatiu FEDER Catalunya 2014-2020.

